# Inhibition of acetyl-CoA carboxylase by spirotetramat causes lipid depletion and surface coat deficiency in nematodes

**DOI:** 10.1101/278093

**Authors:** Philipp Gutbrod, Katharina Gutbrod, Ralf Nauen, Abdelnaser Elashry, Shahid Siddique, Jürgen Benting, Peter Dörmann, Florian M.W. Grundler

## Abstract

Plant-parasitic nematodes pose a significant threat to agriculture causing annual yield losses worth more than 100 billion US$. Nematode control often involves the use of nematicides, but many of them including non-selective fumigants have been phased out, particularly due to ecotoxicological concerns. Thus new control strategies are urgently needed. Spirotetramat (SPT) is used as phloem-moble systemic insecticide targeting acetyl-CoA carboxylase (ACC) of pest insects and mites upon foliar application. Our studies revealed that SPT known to be activated *in planta* to SPT-enol acts as a developmental inhibitor of the free-living nematode *Caenorhabditis elegans* and the plant-parasitic nematode *Heterodera schachtii*. Exposure to SPT-enol leads to larval arrest and disruption of the life cycle. Furthermore, SPT-enol inhibits nematode ACC activity, affects storage lipids, fatty acid composition and disrupts surface coat synthesis. Silencing of *H. schachtii ACC* by RNAi induced similar phenotypes and thus mimics the effects of SPT-enol, supporting the conclusion that SPT-enol acts on nematodes by inhibiting ACC. Our studies demonstrated that the inhibition of *de novo* lipid biosynthesis by interfering with nematode ACC is a new nematicidal mode of action addressed by spirotetramat, a well-known systemic insecticide for sucking pest control.

## Introduction

Nematodes are widespread organisms with diverse lifestyles including free-living microbe feeders, predators, and obligate parasites of plants and animals. Although they employ fundamentally different life-history strategies, many developmental characteristics are shared. Nematodes hatch from an egg and under favorable conditions develop through consecutive molts into fertile adults, which can propagate and repeat their lifecycles. *Caenorhabditis elegans*, the best-studied nematode model organism, is a free-living ubiquitous soil organism. Plant-parasitic nematodes like root-knot and cyst nematodes, on the other hand, have a significant impact on agricultural yields causing a global annual damage of more than 100 billion US$ (Sasser *et al*., 1986). However, little is known about their basic biology. *Heterodera schachtii* a cyst nematode, represents an economically important biotrophic parasite of sugar beet, other Chenopodiaceae and Brassicaceae including the model plant *Arabidopsis thaliana*. Following root invasion, *H. schachtii* induces a feeding site containing a syncytium of cells in the root central cylinder. *H. schachtii* actively feeds from the syncytium that provides all nutrients required for its development. While the nematode develops and grows, the feeding site also enlarges and forms strong sink diverting nutrients away from the plant and providing it to the nematode (Sijmons *et al.*, 1991; Golinowski *et al.*, 1996). After hatching from an egg, vermiform L1 of *C. elegans* and J2 of *H. schachtii* will not develop unless they are supplied with a food source. In *C. elegans*, nutrients derived from feeding on bacteria activate a signaling cascade that promotes its development (Ashraf, 2007). Similarly, in *H. schachtii* the development of J2s is initiated after feeding from the previously induced syncytium. This indicates that in both nematodes, developmental initiation is tightly linked to nutrient-dependent signaling. While feeding, cyst nematodes secrete a hydrophobic surface coat (subcrystalline layer) onto their epicuticle that is likely involved in the protection against environmental stress and in the interaction with other organisms (Brown *et al*., 1971; Zunke, 1986; Endo and Wyss, 1992; Davis and Curtis, 2011).

Control of plant-parasitic nematodes is challenging and often requires complex integrated pest management practices that also involve the use of synthetic nematicidal compounds (Haydock *et al*., 2013). However, chemical control of nematodes is seen controversial by the public due to the potential damage caused to the environment and non-target organisms. Nevertheless, the search for new nematicides is ongoing (Faske and Hurd, 2015; Vang *et al*., 2016). In order to predict any potential risks associated with the use of nematicides it is of essential importance to understand their mode of action and to study their exerted effects.

Spirotetramat (SPT) belongs to the class of cyclic keto-enols and is used as a systemic insecticide in crop protection. Within the plant, SPT is hydrolyzed to the corresponding SPT-enol, the presumed active form of SPT. SPT-enol is transported via the phloem, and therefore distributed throughout the plant. In insects and mites, SPT-enol suppresses fatty acid biosynthesis by inhibiting the action of acetyl-CoA carboxylase (ACC). More specifically, SPT-enol inhibits the carboxyltransferase partial reaction and shows a competitive mode of inhibition with respect to acetyl-CoA and is uncompetitive with respect to ATP (Nauen *et al.*, 2008; Brück *et al.*, 2009; Lümmen *et al.*, 2014). Target identification in insects and mites is further supported by studies on the structurally related keto-enol spiromesifen (Karatolos *et al.*, 2012) suggesting a common mode of action.

ACC catalyzes the initial and rate-limiting step in fatty acid *de novo* synthesis synthesizing malonyl-CoA via the carboxylation of acetyl-CoA. Malonyl-CoA is used as a substrate for *de novo* fatty acid biosynthesis by fatty acid synthase (FAS). Fatty acids produced by FAS may be further elongated by the ER-localized acyl-CoA elongation enzymes that also require malonyl-CoA. Fatty acids may be desaturated and are eventually assembled into complex lipids. Lipids are vital constituents of cells as they are involved in membrane biogenesis (e.g. phospholipids), energy storage (triacylglycerol) and modulation of growth and development (e.g. glycosylceramides) (Watts *et al.*, 2002; Barber *et al.*, 2005; Zhang *et al.*, 2011; Zhu *et al.*, 2013). ACC activity is highly responsive to the overall energy status of a cell which is in turn regulated by nutritional cues. The signal transduction depends on AMP-activated protein kinase (AMPK) that in times of fasting (high AMP, low ATP, low insulin) keeps ACC phosphorylated and thus inactive. In times of abundant nutrients and a high energy charge (low AMP, high ATP, high insulin) AMPK becomes inactive and the proportion of dephosphorylated ACC increases and more malonyl-CoA is synthesized (Berg *et al.*, 2002).

ACC activity is crucial for embryonic and post-embryonic development of various organisms highlighting the importance of fatty acid *de novo* synthesis for development. In line with these findings, developmental defects induced by SPT-enol in insects and mites include incomplete molting and reduced fecundity. Furthermore, inhibitory effects were also described for nematodes (Smiley *et al*., 2011; Vang *et al*., 2016). However, the underlying mode of action of SPT on nematodes has not been studied. Therefore, we set out for studying the effects and to elucidate the mode of action of SPT on *C. elegans* and *H. schachtii*, i.e. on free-living and parasitic nematodes, both under developing and non-developing conditions.

## Results

### SPT-enol is not acutely toxic for nematodes

To test for an acute toxic effect of SPT-enol we examined its impact on non-developing stages of nematodes, i.e. *C. elegans* starved L1 and *H. schachtii* J2 larvae. When incubated with SPT-enol, the two nematode species were not differently inactivated during the following time course analysis as compared to the solvent control (Supplemental Figure 1A). When the SPT-enol was removed by washing and nematodes were allowed to feed and grow, larval fate was unaffected (Figure 1C, Supplemental Figure 1B).

**Figure 1:**
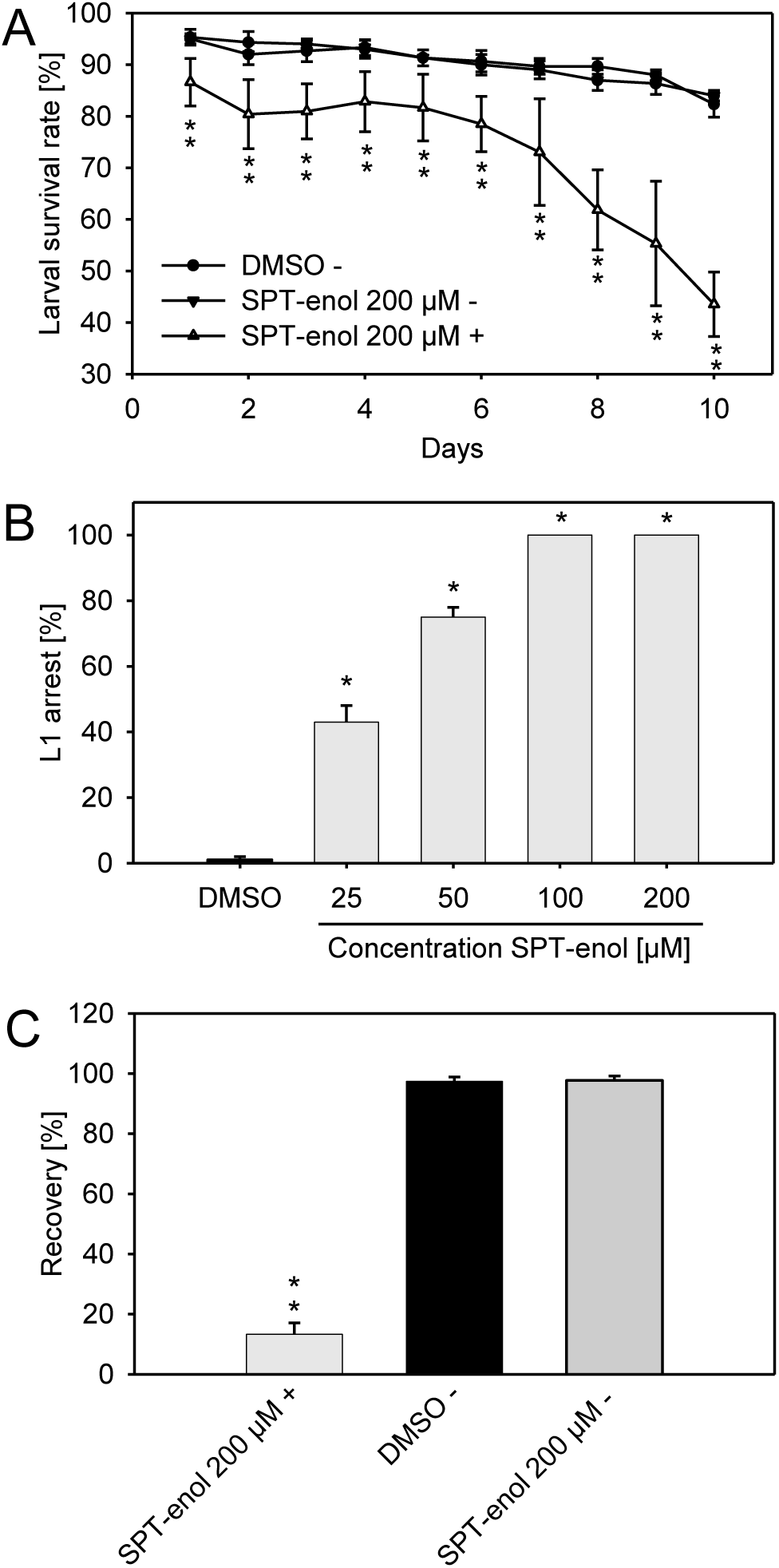
Effect of SPT-enol on *C. elegans* development. (A) Time course experiment on survival rate of non-developing and arrested *C. elegans* nematodes. The data are given in average ± SD (n=3). For each time point, Student’s t-test was performed for SPT-enol+ against DMSO (control) and SPT-enol+ against SPT-enol– (*P < 0.001). (B) Concentration-dependent effect of SPT-enol on *C. elegans* larval arrest. The data are given in average ± SD (n=3). For each concentration of SPT-enol, Student’s t-test was performed for SPT-enol against DMSO (control) (*P < 0.001). (C) Recovery of non-developing L1 larvae and SPT-enol-arrested larvae of *C. elegans*. The data are given in average ± SD (n=3). Student’s t-test was performed for SPT-enol 200 µM + against DMSO (control) and SPT-enol 200 µM + against SPT-enol 200 µM -(*P < 0.001). In these experiments, (+) and (–) indicate presence or absence of a food source. DMSO plus food source was excluded from the analysis since larvae develop into adults under these conditions.

To further induce SPT-enol uptake by *H. schachtii* J2 we stimulated them with octopamine, which has been shown to enhance xenobiotic transfer from the surrounding medium into the nematode (Urwin *et al.*, 2002). Similar to our previous observations, preincubation with octopamine and SPT-enol did not affect nematode development (Supplemental Figure 1C). The findings that SPT-enol neither inactivated nematodes over time nor altered their developmental fate after incubation suggest that SPT-enol is not acutely toxic for larval stages that have not yet initiated development.

### SPT-enol inhibits nematode development

Next, we tested if SPT-enol would affect the two nematode species during their development, which starts as soon as nutrients are available. SPT-enol inhibited *C. elegans* development in a concentration dependent manner, causing early arrest at the L1 stage. At a concentration of 100 μM or above L1 arrest occurs uniformly (Figure 1B). In a time course experiment, those early arrested larvae became inactive over time (Figure 1A). To asses if the arrest was reversible, arrested L1 were washed after 2 days and transferred to nematode growth medium (NGM) plates without SPT-enol. Approximately 10% of those arrested L1 were able to recover and became adults whereas all L1 incubated without food source were able to develop irrespective of the presence of SPT-enol (Figure 1C).

To study the effect of SPT on developing *H. schachtii*, a host plant, in this case *A. thaliana* is required. Therefore, we first analyzed if SPT is hydrolyzed to SPT-enol and transported basipetally from the leaf to the root. In addition, we addressed the question if this treatment would affect the growth of *A. thaliana*, to exclude phytotoxicity as a possible cause of altered nematode development.

After SPT had been applied onto leaves of *A. thaliana*, SPT-enol could be detected in the roots and in the growth medium of the plants (Supplemental Figure 2). SPT applied at a concentration of 200 mM did not affect shoot and root growth (as determined by measuring fresh weight) of the treated plants (Supplemental Figure 3). Taken together these results suggested that SPT is hydrolyzed to SPT-enol *in planta* and translocated basipetally (leaf-to-root) without causing an apparent harm to the plant. Next, we assessed the development of *H. schachtii* infecting SPT-treated plants. Similar to the effect of SPT-enol on *C. elegans*, development of *H. schachtii* infecting SPT-treated plants was arrested in early stages in a concentration-dependent manner (Figure 2A). In addition, females that did develop on SPT-treated plants were significantly smaller than control nematodes (Figure 2B). Interestingly, the feeding sites of these developmentally suppressed females were not significantly smaller than those of control nematodes, indicating that the plant response to the infection was not affected. In contrast, feeding sites of nematodes arrested at an early stage were significantly smaller (Figure 2A, Supplemental Figure 4). Since the feeding sites were induced and nematodes started development, the effect of SPT-enol seems to be restricted to nematodes that have initiated development. The data suggest that SPT-enol acts as a developmental inhibitor with a partially reversible effect. Additionally, the juvenile arrest seen in *H. schachtii* seems to be a direct effect of SPT-enol on the nematode rather than indirectly through phytotoxicity.

**Figure 2:**
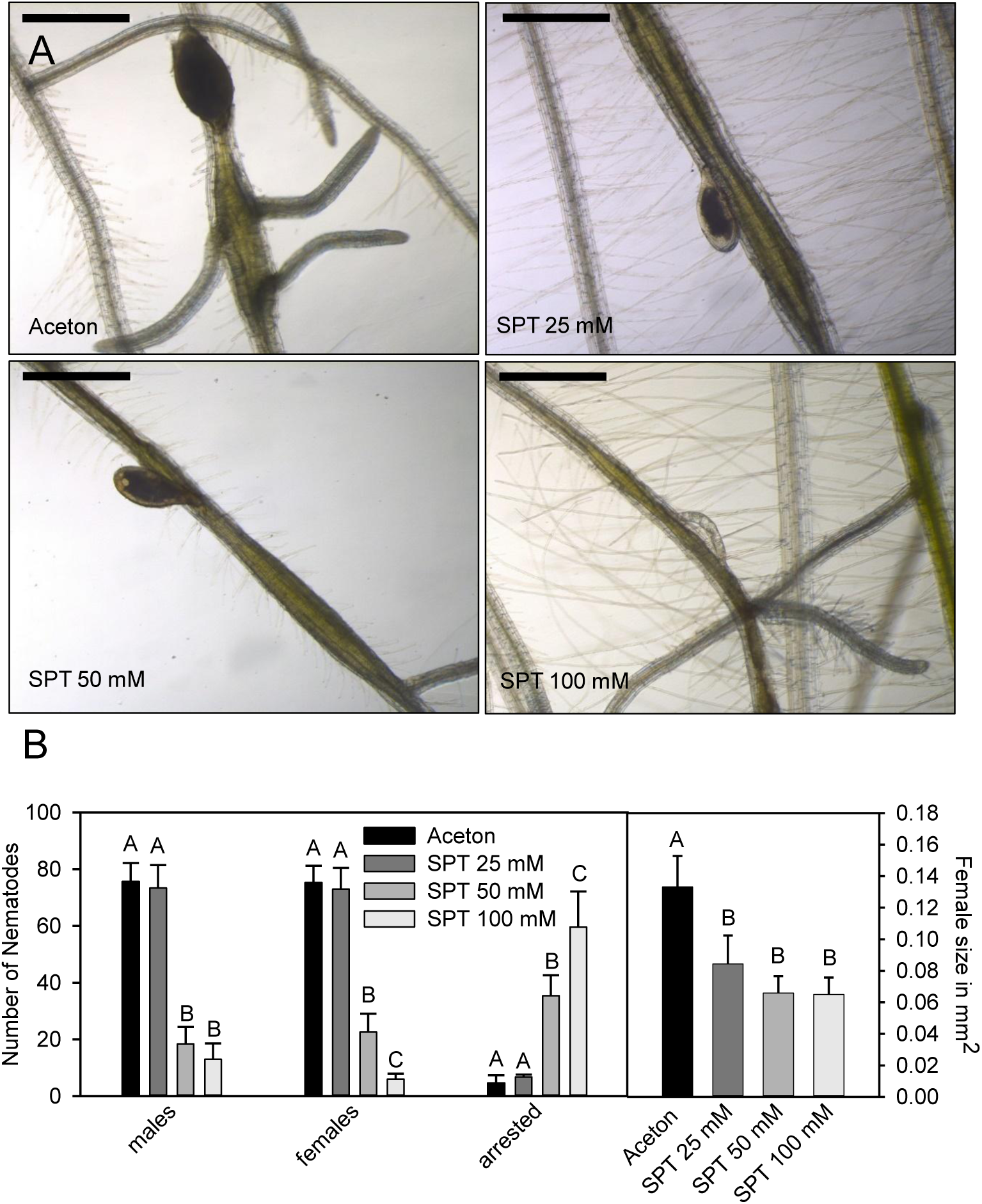
Concentration-dependent effect of foliar application of SPT on *H. schachtii* development. (A) Representative images of *H. schachtii* females in interaction with *A. thaliana* roots after the leaves of the respective plants have been treated with 25 mM, 50 mM and 100 mM SPT. The black bar represents 500 µm. (B) Concentration-dependent effect of foliar SPT application on *H. schachtii*. The data are given in average ± SD. Anova on ranks for males P = 0.002, Tukey Test, N=10; Anova on ranks for females P < 0.001; Tukey Test, N=10; Anova on ranks for arrested P = 0.001, Tukey Test N=10; Anova for sizes P < 0.001, Tukey Test, N=30. Please note that males, females, arrested nematodes and female sizes were analyzed for statistical significance separately.

### SPT-enol interferes with lipid metabolism in nematodes by inhibiting ACC

SPT-enol was shown to interfere with lipid metabolism in insects and mites (Nauen *et al.*, 2008; Brück *et al.*, 2009). Given the importance of lipids for development we investigated the effect of SPT-enol on *C. elegans* storage lipids. When treated with SPT-enol, early arrested L1 larvae were found to be devoid of intestinal lipid droplets as shown by nile red staining. In contrast, lipid droplets of L1 incubated for the same time without food source showed similar lipid droplet staining irrespective if SPT-enol was present or not (Figure 3).

**Figure 3:**
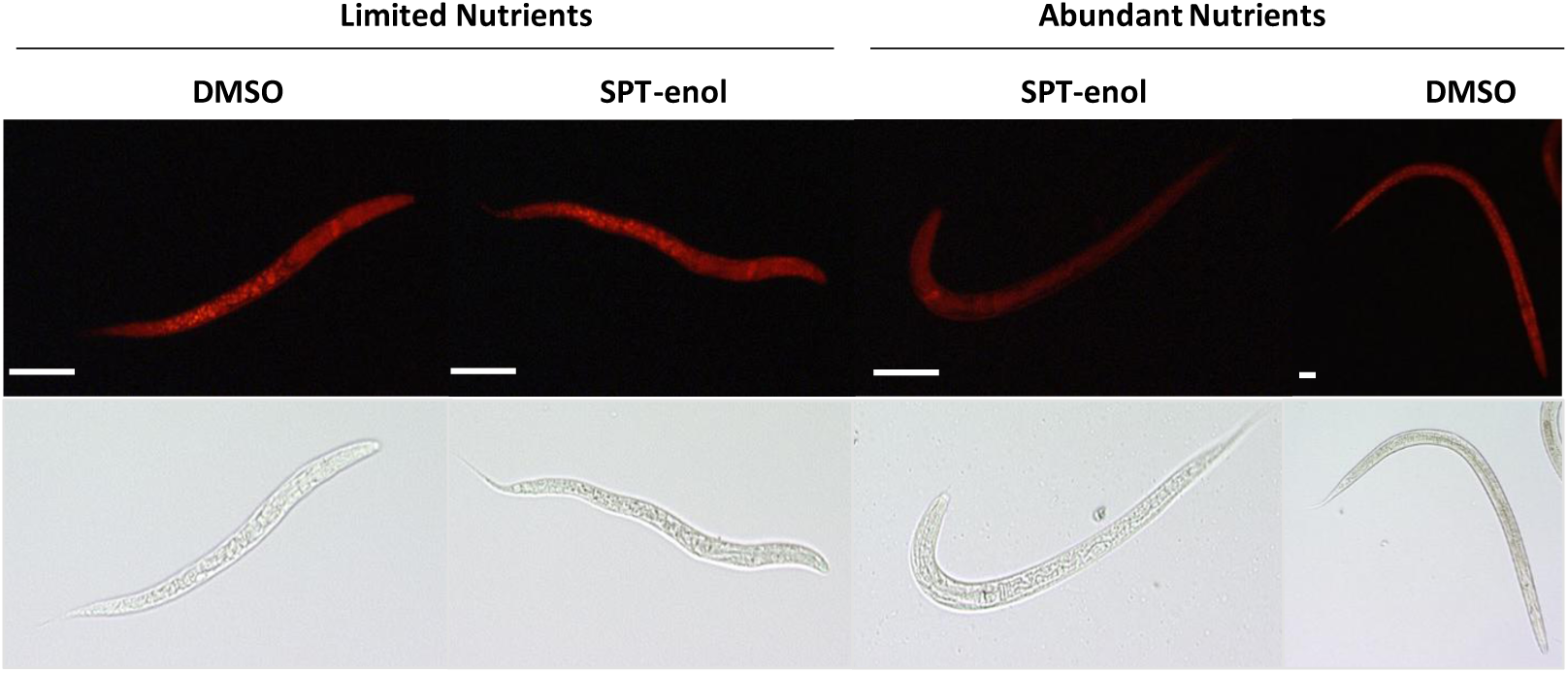
Nile red staining of intestinal lipid droplets of *C. elegans*. Synchronized L1 were incubated in liquid S-medium with 200 µM SPT or DMSO (control) with and without nutrient source. After 48 hours, the nematodes were fixed and stained with nile red and the number of lipid-depleted nematodes was counted. Only nematodes that were incubated with SPT-enol in the presence of a food source show intestinal lipid droplet depletion (92.3 ± 2.5% of total nematodes, average and SD, n=3). Nematodes supplied with food and DMSO start to develop. The white bar represents 50 µm.

Absence of intestinal lipid droplets maybe associated with an altered fatty acid metabolism. Therefore, we analyzed total fatty acids in SPT-enol treated nematodes. We found a reduction of C18 fatty acids while C16 fatty acids were relatively increased (Figure 4A). To get a more detailed view on storage lipid changes, we determined the triacylglycerol (TAG) content and composition using direct infusion Q-TOF mass spectrometry. We found that SPT-enol treated nematodes had a significantly lower total TAG content as compared to the solvent control (Figure 4B). The reduction of C18 observed in total fatty acid composition is reflected in changes in TAG composition. When analyzing the most abundant TAG molecular species it became apparent that 18:1-containing TAG molecules were reduced while TAGs containing 16:1-and/or 17:cyclo-fatty acids exclusively were unaffected (Figure 4C). Together this indicates that SPT-enol affects the nematode’s storage lipids, by specifically reducing C18-containing TAGs, linking developmental inhibition to fatty acid metabolism. In nematodes, 18:1 is produced via elongation and desaturation of 16:0, while 16:0 can be either taken up by the diet or produced by *de novo* fatty acid synthesis. Both fatty acid elongation and *de novo* synthesis require malonyl-CoA. The sole route for malonyl-CoA production in nematodes is carboxylation of acetyl-CoA by ACC. Therefore, we tested wether SPT-enol is able to inhibit ACC activity *in vitro*. Enzyme assays using radiolabeled sodium bicarbonate showed that *C. elegans* ACC activity is inhibited by SPT-enol with an IC_50_ of 50 μM (Figure 4D). This supports the conclusion that the altered fatty acid metabolism in *C. elegans* is linked to ACC inhibition.

**Figure 4:**
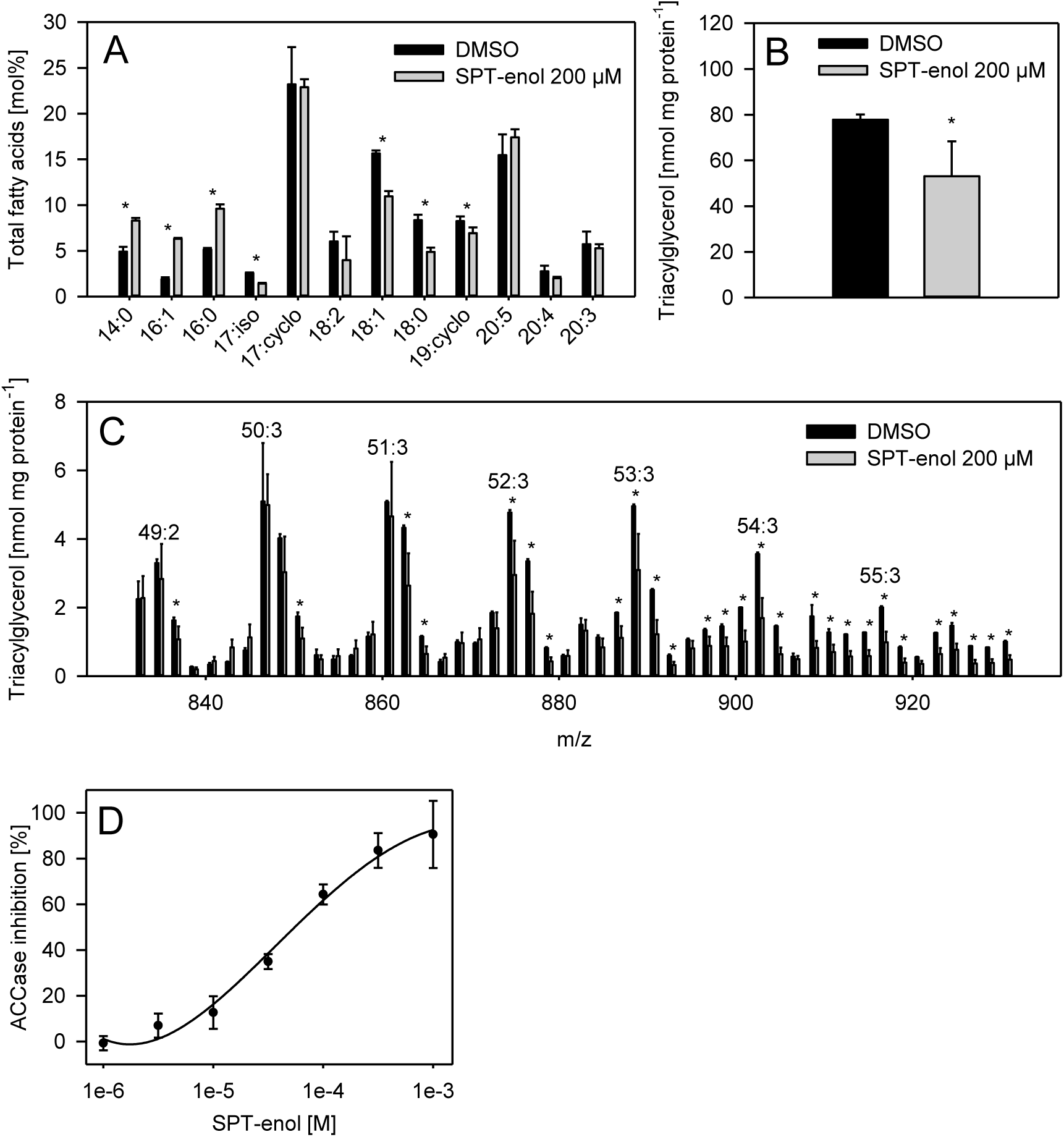
Effect of SPT-enol on fatty acid composition, TAG content and ACC activity of *C. elegans.* (A) Total fatty acid composition of *C. elegans* (mol%) after treatment with DMSO (control) or with 200 µM SPT-enol. The data are given in average ± SD (n=3). Student’s t-test was performed (*P < 0.05). (B) Total TAG content of *C. elegans* (nmol mg protein-1) after treatment with DMSO (control) or with 200 µM SPT-enol. The data are given in average ± SD (n=3). Student’s t-test was performed (*P < 0.05). Representative TAG molecular species are shown in supplemental table 1 (C) Molecular species of TAGs of *C. elegans* (nmol mg protein-1) after treatment with DMSO (control) or with 200 µM SPT-enol. The data are given in average ± SD (n=3). Student’s t-test was performed (*P < 0.05). (D) Relative inhibition of *C. elegans* ACC activity (%) by different concentrations of SPT-enol. The data are given in average ± SD (n=5).

Our data show that developmental inhibition observed in both nematode species after SPT-enol treatment is the result of dysfunctional lipid metabolism due to specific inhibition of ACC by SPT-enol.

### RNAi mediated silencing of nematode ACC mimics the effect of spirotetramat

To elucidate whether the two nematode species are subject to a common mode of action of SPT-enol on ACC, we searched for the ACC sequence in *H. schachtii*. Using a *H. schachtii* transcriptome data base and the *C. elegans* ACC (POD-2) sequence as query, we retrieved a single contig with a translated ORF of 2024 amino acids. The predicted protein contains all functional domains required for ACC activity and is similar to other nematode ACCs (Supplemental figure 5A, 5B). Hence, this implies that we found the *H. schachtii* ortholog of the *C. elegans* ACC (POD-2) and therefore named it *Hs*ACC (HsPOD-2).

To test if *Hs*ACC is relevant for the development of *H. schachtii* we investigated the effect of its knock-down upon RNAi. Soaking of J2 in a solution of dsRNA targeted against *Hs*ACC reduced transcript levels by 90% (Figure 5B). When allowed to develop on *A. thaliana*, the numbers of males and females were not significantly different, but the developed females were significantly smaller as compared to the control (Figure 5A, 5C). To check if transcript reduction was persistent we quantified *Hs*ACC transcript levels in the developed females 10 days after dsRNA treatment. However, at this stage, *Hs*ACC transcript levels were no longer significantly different between control and treated females (Figure 5B).

This suggests that the *Hs*ACC transcript is involved in *H. schachtii* development. Moreover, its transient knock-down phenocopies the suppressive effect of SPT-enol observed at concentrations of 25 mM. Taken together, we conclude that SPT-enol acts on a common nematode target, which is ACC.

### ACC inhibition disrupts surface coat formation

We also observed that ACC inhibition caused by either foliar SPT application or soaking in dsRNA targeting *Hs*ACC caused a “smoothening” of the cyst nematodes surface indicating a defect of the hydrophobic surface coat (subcrystalline layer). To visualize hydrophobic structures on the nematodes surface, plates were stained with nile red. It was observed that both treatments (SPT and dsRNA) lead to a reduced surface staining (Figure 6A). The surface coat of other cyst nematodes has been shown to contain very long chain fatty acids (VLCFA) (24:0 and 26:0) and their corresponding calcium salts (Brown *et al*., 1971). Indeed, surface lipid extracts (by chloroform-dipping) of female nematodes contained VLCFA (mainly 24:0 and 26:0 but also 22:0) confirming and extending previous results (Supplemental figure 6).

**Figure 6:**
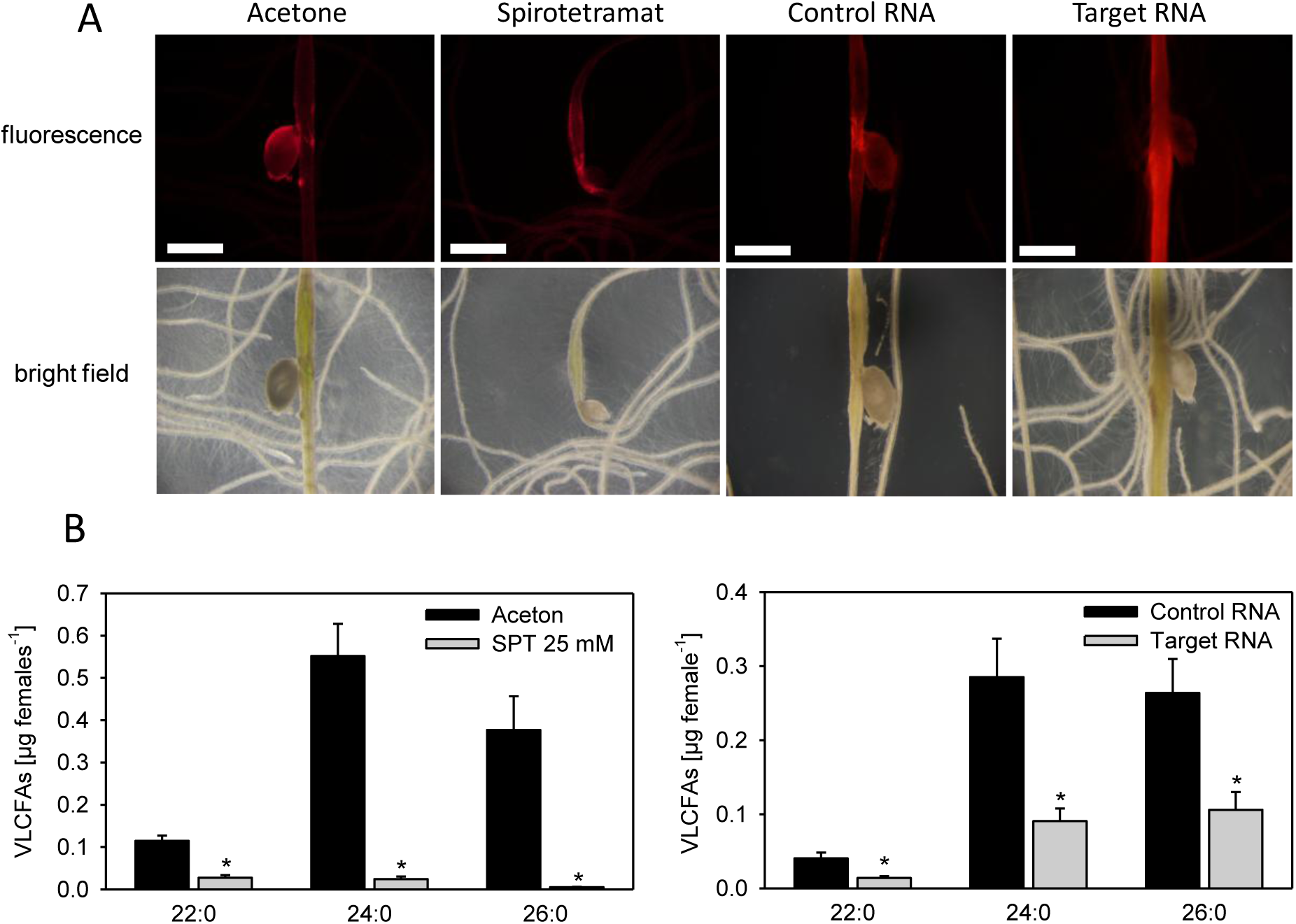
Effect of ACC inhibition on surface lipids of *H. schachtii*. (A) Representative images of nile red-stained surface coats of *H. schachtii* females after foliar SPT application and after RNAi against HsACC. The bar represents 500 µm. (B) VLCFA content of female *H. schachtii* after foliar SPT application and RNAi against HsACC. The data are given in average ± SE (n=8). Student’s t-test was performed (*P < 0.05)

Quantification of VLCFA after both treatments (foliar SPT application and RNAi against *Hs*ACC) revealed a reduction of these fatty acids as compared to their corresponding controls (Figure 6B). This argues for VLCFA as an essential component of the cyst nematodes surface coat and that ACC inhibition disrupts its formation.

## Discussion

In this article we show that SPT-enol interferes with the development of nematodes by inhibiting ACC. SPT-enol affects the development of two distantly related nematode species, causes lipid droplet depletion and impairs malonyl-CoA dependent fatty acid synthesis. Enzyme inhibition experiments confirmed ACC as the main target of SPT-enol. Furthermore down-regulation of its expression by RNAi mimics the effect of SPT-enol in both *C. elegans* and *H. schachtii*. Together, this strongly supports the scenario that ACC is the target of SPT-enol in nematodes.

ACC catalyzes the initial step in fatty acid biosynthesis converting acetyl-CoA to malonyl-CoA. Malonyl-CoA serves multiple purposes. It supplies C2 units for fatty acid *de novo* synthesis by fatty acid synthase (FAS), is required for fatty acid elongation and regulates mitochondrial β-oxidation (Watts, 2008). Fatty acids are constituents of structural components of membranes (e.g. phospholipids, sphingolipids), energy stores (triacylglycerol) and are important modulators of development (e.g. phosphoinositides, glycosylceramides) (Watts 2008; Zhu *et al.*, 2014).

In agreement with the important functions of fatty acids, ACC inactivation was found to be lethal in various organisms including nematodes, flies, mice and yeast (Hasslacher *et al.*, 1993; Tagawa *et al.*, 2001; Abu-Elheiga *et al.*, 2005; Parvy *et al.*, 2012). Also, polyunsaturated fatty acid (PUFA) synthesis is crucial for development beyond the L1 stage but can be complemented by dietary fatty acids (Brock *et al.*, 2007). In line with these observations, post embryonic knock-down of *C. elegans* POD-2 encoding ACC led to larval arrest (Li *et al.*, 2011). However, we were not able to rescue the SPT-enol induced developmental arrest in nematodes by dietary supplementation. Similarly ACC defective yeast could also not be rescued by exogenous supply of lipids (Hasslacher *et al.*, 1993). The inability to rescue ACC deficiency pinpointed to a defect in supplying malonyl-CoA for very long chain fatty acid (VLCFA) production which is in turn required for the synthesis of sphingolipids (Tehlivets *et al.*, 2007). Indeed, for *C. elegans* glycosylceramide synthesis is essential for the establishment of cell-polarity and is also required for the activation of TORc signaling, a key activator of development (Zhang *et al.*, 2011; Zhu *et al.*, 2013). Furthermore, the inability to synthesize glycosylceramides has been linked to L1 arrest in *C. elegans (*Marza *et al., 2009*). Therefore, the lack of fatty acid *de-novo* synthesis inhibits the synthesis of specific lipids that are essential for development.

SPT-enol-treated *C. elegans* had lower TAG levels and lacked intestinal lipid droplets when nutrients were present. In addition to its important role in fatty acid synthesis, malonyl-CoA inhibits the action of carnitine palmitoyl transferase (CPT) thereby preventing mitochondrial β-oxidation (Stryer *et al.*, 2002). The absence of malonyl-CoA due to ACC inhibition may explain intestinal lipid droplet depletion under high nutrient conditions. In contrast, lipid droplets in non-developing L1 appeared similar irrespective of the presence of SPT-enol. Interestingly, lowering ACC expression levels by RNAi increases hepatic fat oxidation specifically in the fed state (Savage *et al.*, 2006). In support of these findings, knock-down of ACC by RNAi in feeding *C. elegans* also led to lower storage lipid (TAG) content (Li *et al.*, 2011). We also observed the inactivation of arrested and lipid depleted *C. elegans* L1 larvae over time. This inactivation may be associated with increased cell death that had been reported in lipid depleted cancer cells due to ACC inhibition (Becker *et al*., 2007). Thus, lipid depletion seems to drive nematode inactivation possibly by increasing cell death.

Support for a common target of SPT-enol in nematodes is provided by our studies on plant-parasitic *H. schachtii*. SPT does apparently not affect *A. thaliana* growth, SPT-enol is phloem-mobile and the nematodes are arrested in development after feeding site induction. This suggests that SPT acts directly on developing *H. schachtii* rather than through a phytotoxic effect. Both nematode species actively transcribe an ACC mRNA and silencing of *Hs*ACC by RNAi phenocopies the suppressive effect of SPT-enol on female development. This effect indicates a role for *Hs*ACC also in *H. schachtii* development and further supports target identification for nematodes.

The IC_50_ of SPT-enol for insect and mite ACCs *in vitro* is in the nM range and thus approximately 1000-fold lower than for nematode ACC (Nauen *et al.*, 2008; Brück *et al.*, 2009; Lümmen et al., 2014). In line with this result, *in vivo* inhibition of nematode development requires relatively high concentrations of SPT-enol as compared to insects and mites. Therefore, the target affinity of SPT-enol for nematode ACC appears low. This may not be surprising since SPT was developed as an insecticide and the phylogenetic distance and hence ACC sequence differences between nematodes and insects are relatively large. Nevertheless, a formulation of SPT has been reported to reduce cyst nematode populations in wheat (Smiley *et al.*, 2011). This indicates that factors other than the target affinity may ultimately determine the efficacy in the field.

Interestingly, we also found that ACC inhibition disrupts surface coat formation. The surface coat of cyst nematodes (sub-crystalline layer) contains VLCFA and is likely serving multiple roles. It provides a physical barrier protecting the nematode against environmental stress and is involved in the interaction with other organisms (Brown *et al*., 1971; Davis and Curtis, 2011). A lack of malonyl-CoA due to ACC inhibition would abolish VLCFA synthesis thereby disrupting surface coat formation. Therefore, the control of nematodes in the field may have also been the result of indirect effects due to loss of protective barriers or altered biotic interactions.

We found that SPT-enol is not acutely toxic for nematodes but suppresses their development. Non-developing larval stages of *H. schachtii* and *C. elegans* are not inactivated by SPT-enol and their developmental fate is unaffected following SPT-enol removal. Interestingly, SPT-enol inhibition of ACC was shown to be uncompetitive with respect to ATP suggesting that it binds to the enzyme-substrate complex (Lümmen *et al.*, 2014). Since ATP content is low under non-developing conditions due to lack of nutrients, SPT-enol may not be able to efficiently bind to the ACC complex. In contrast, under fed conditions ATP is high and also the effect of an uncompetitive inhibition increases. Thus, the inability to provoke effects in non-developing nematodes may be the result of insufficient ATP levels which are required for SPT-enol to inhibit ACC activity.

In conclusion, this is the first report of ACC as a pharmacological target for nematode control. The findings reported here have significant implications for the treatment of parasitic nematodes in agriculture and may also be translated into novel strategies to fight animal-parasitic nematodes. Additionally, the discovery of specific small molecule inhibitors is an important complement for basic research, especially where sophisticated molecular tools are unavailable.

## Materials and Methods

### Strains

*C. elegans* Bristol N-2 and *E. coli* OP50 were kindly provided by Prof. Schierenberg, University of Cologne. *C. elegans* was maintained at 20°C using standard methods and synchronous L1 stages were obtained using alkaline hypochlorite treatment and hatching in M9 (Stirnagle *et al.*, 2006).

### Drug preparation

For experiments with nematodes in solution, SPT-enol was dissolved in DMSO. For foliar application, SPT was dissolved in acetone.

### Liquid culture

Liquid cultures were set up in 24 well plates using 890 µL S-complete, 10 µL active ingredient in DMSO (1% final concentration) and 100 µL M9 buffer containing 250-300 L1s. For starvation experiments bacteria were omitted from the S-complete medium. Liquid cultures were maintained on a shaker with 180 rpm at 20°C.

### Drug recovery assay

Nematodes were collected from liquid cultures, washed twice with M9 and transferred to ngm plates containing *E.coli* OP50. After 3 days developed nematodes were counted.

### Nematode fixation and Nile red staining

*C. elegans* were fixed and stained as described in Barros *et al*. (2012). For visualization of *H. schachtii* surface lipids, plates were overlayed with 2 mL of a solution of 2 µg/mL Nile red and incubated overnight.

### Imaging

Imaging was performed with a binocular microscope (Leica KL200 LED) or with a binocular microscope (Olympus SZX16) equipped with suitable filters to observe Nile red fluorescence.

### Lipid quantification

Lipids were extracted from 50 µL mixed-stage worm pellet cultured in S-complete medium with 200 µM SPT-enol or DMSO as control. Triacylglycerol was purified and analyzed as described previously (vom Dorp *et al.*, 2013). Additionally, isolated lipids were transmethylated with 1 N HCl in methanol for 20 min at 80°C. Fatty acid methyl esters (FAMEs) were analyzed by gas chromatography (Agilent 7890 GC device) coupled to mass spectrometry (GC-MS). FAMEs of different chain lengths and degrees of unsaturation were separated on an HP-5MS column using a gradient of 70°C to 310°C at 10°C/min, followed by a hold at 310°C for 1 min, afterwards the temperature decreases to 70°C again. Quantification of fatty acids was based on the comparison of the peak area to that of an internal standard, pentadecanoic acid (15:0), which was added in a specified amount (5 µg).

### Fatty acid composition of surface lipids

To determine the very long chain fatty acid composition of surface lipids, five females 14 days old were picked from a plate and incubated in chloroform for 1 min. The chloroform extract was transferred to a fresh glass tube, the organic solvent evaporated and the lipids were transmethylated with 1 N HCl in methanol for 20 min at 80°C. For quantification of very long chain fatty acids, single females were picked from a plate transferred to a fresh glass tube and transmethylated with 1 N HCl in methanol for 2 hours at 80°C. Analysis with GC-MS was performed as described above.

### Acetyl-CoA Carboxylase assay

Total proteins were extracted from 3 mL of dense *C. elegans* pellet using a protein isolation buffer containing 20 mM HEPES, 150 mM NaCl, 1 mM EDTA, 1 mM DTT, 10% glycerol, 1% protease inhibitor cocktail "Halt” (Thermo Scientific). Determination of ACC activity and inhibition by SPT-enol was performed as described in Lümmen *et al*. (2014).

### Residue Analysis

10-day-old *A. thaliana* Col-0 plants cultured on 0.2 KNOP plates were treated with 3 μL 200 mM SPT as described above. After one, three and 10 days, roots and agar were sampled. Roots were thoroughly rinsed with H_2_0, blotted dry, weighed, frozen in liquid N_2_ and crushed in a Precellys homogenizer (Peqlab Biotechnology). Residues were extracted with 1 mL acetonitrile/H_2_0 (9:1). Quantification was performed according to Schöning (2008).

### Toxicology tests for *H. schachtii*

*H. schachtii* was incubated in a solution containing M9 buffer and 500 μM SPT-enol. The final DMSO concentration was 1%. Non-moving nematodes were counted daily. After 48 h, nematodes in J2 stage were washed with M9 buffer and transferred to 0.2 KNOP plates containing 10-day-old *A. thaliana*.

### *H. schachtii* infection assay

Infection assays were performed on 0.2 KNOP medium as described in Simmons *et al*. (1991). Per single experiment, 5 plants per petri dish were used. The total numbers of female and male nematodes per plate were counted at 10 DPI and females and feeding site sizes were determined at 14 DPI. Just before nematode inoculation 3 μL spirotetramat in acetone with the indicated concentrations were applied foliarly onto the largest green leaf. Acetone was allowed to evaporate.

### Acetyl-CoA carboxylase transcript identification

A transcript sequence containing a full-length ORF of *H. schachtii* acetyl-CoA carboxylase was identified in a transcriptome database based on sequence similarity to *C. elegans* POD-2 that encodes acetyl-CoA carboxylase. Functional domains were predicted using Pfam (pfam.xfam.org). In order to compare the positions of the predicted functional domains, the HsACC protein sequence was aligned against the *C. elegans* POD-2 sequence (CLC genomics workbench version 6.0).

### Sequence analysis

Protein sequences corresponding to full ORF orthologs of *C. elegans* POD-2 were collected from wormbase parasite (parasite.wormbase.org). Domain structures of ACC of *C. elegans* and *H. schachtii* were illustrated using DOG 2.0 (Ren et al. 2009). Sequences were aligned using ClustalW. The evolutionary history was inferred using the Neighbor-Joining method (Saitou *et al*., 1987). The optimal tree with the sum of branch length = 4.54642297 is shown. The percentage of replicate trees in which the associated taxa clustered together in the bootstrap test (1000 replicates) are shown next to the branches (Felsenstein, 1985). The tree is drawn to scale, with branch lengths in the same units as those of the evolutionary distances used to infer the phylogenetic tree. The evolutionary distances were computed using the p-distance method (Nei et al. 2000) and are in the units of the number of amino acid differences per site. The analysis involved 60 amino acid sequences. All positions containing gaps and missing data were eliminated. There were a total of 234 positions in the final dataset. Evolutionary analyses were conducted in MEGA6 (Tamura *et al*., 2011).

### dsRNA synthesis

Using GW primers, a GFP fragment was amplified from pMDC107 and an ACC fragment was amplified from *H. schachtii* (J2s) cDNA and both cloned into pDONR207. dsRNA was synthesized *in vitro* with T7 overhang primers listed below using MEGAscript^®^ T7 Transcription Kit (Ambion, Life technologies) according to the manufacturer protocol.

### RNAi of J2s

Induction of RNAi was performed in soaking buffer with 1 μg/μL dsRNA as described by Urwin *et al*. (2002) with modifications from Sukno *et al*. (2007) for transcript quantification by qPCR. dsRNA of a GFP fragment was used as a control RNA.

### Nematode RNA extraction and cDNA synthesis

RNA from nematodes was extracted using the the NucleoSpin RNA Plant Kit (Macherey Nagel) according to the supplier’s recommendations for low amounts of RNA. Residual DNA was digested using Turbo DNase (Life Technologies) according to the manufacturer’s protocol. cDNA was synthesized using the High-Capacity cDNA Reverse Transcription Kit (Applied Biosystems).

### qPCR

qPCR was performed using the Fast SYBR Green Master Mix (Life Technologies) on a StepOnePlus System (Life Technologies). qPCR primers for ACC quantification were validated for efficiency and specificity. Actin was used as the reference gene (Patel *et al.*, 2010). Relative transcript levels were calculated by the comparative CT method (Schmittgen *et al.*, 2008) using the StepOnePlus System Software (Life technologies).

### Statistics

Statistics were performed with Microsoft Office Excel and SigmaPlot.

## Acknowledgement

We would like to thank Peter Lümmen, Claudia Wehr, Gisela Sichtermann and Stefan Neumann for their technical support during the course of this project.

---

**Supplemental Figure 1:**
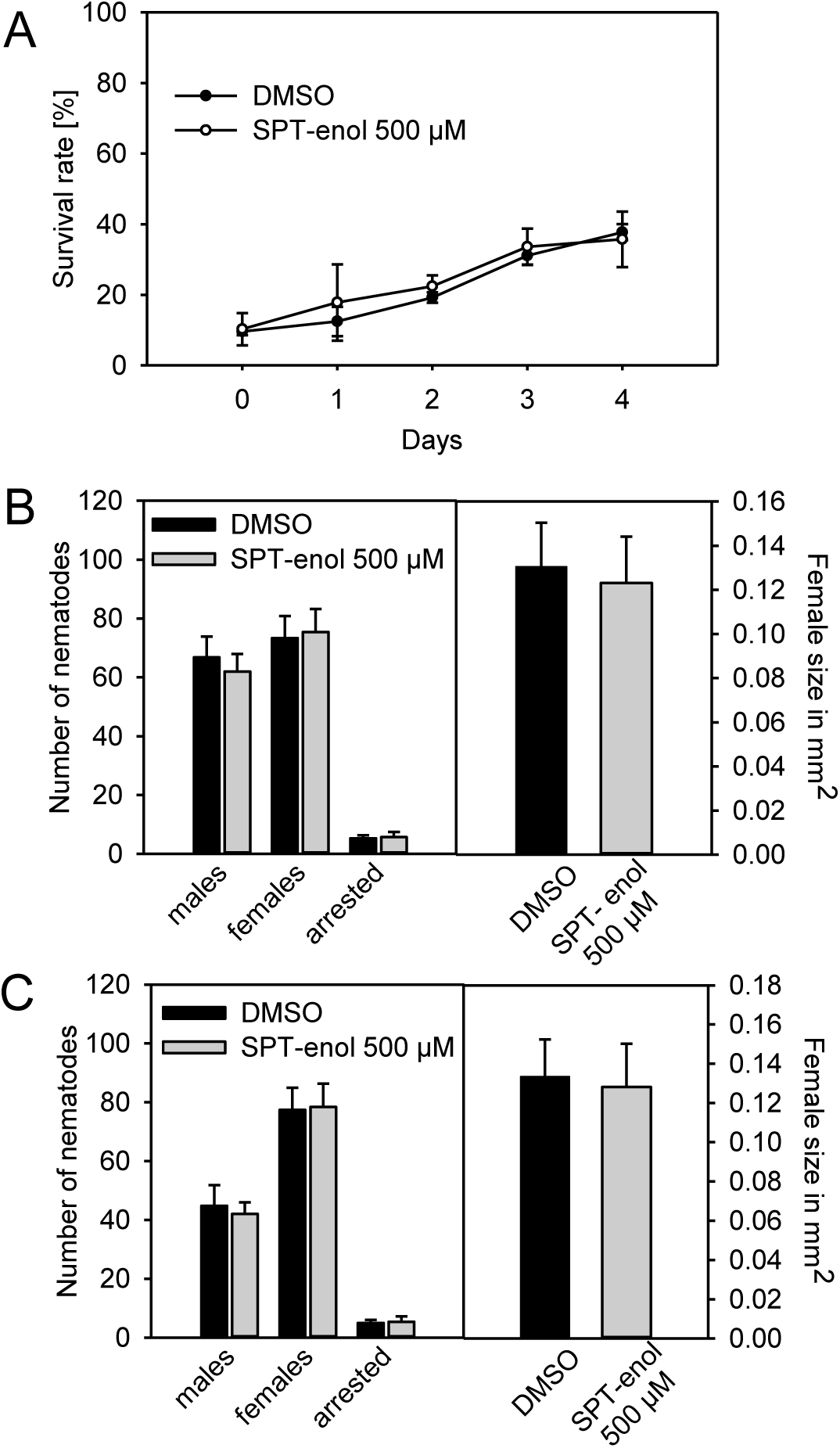
Effect of SPT-enol pre-incubation on J2 mortality and development of H. schachtii. (A) Relative survival rate (%) of *H. schachtii* J2s larvae after treatment with 500 µM SPT-enol. *H. schachtii* J2 larvae were incubated in M9 buffer containing 500 µM SPT-enol or DMSO (control). The number of moving nematodes was counted daily. The data are given in average ± SD (n=5). (B) Number of nematodes and sizes of females after treatment with 500 µM SPT-enol. *H. schachtii* J2 larvae were incubated in M9 buffer containing 500 µM SPT-enol or DMSO (control) and transferred onto plants after 48 hours. The data are given in average ± SD (n=10 for nematode numbers and n=30 for female sizes). (C) Number of nematodes and sizes of females after treatment with 500 µM SPT-enol and 50 mM octopamine. *H. schachtii* J2 larvae were incubated in M9 buffer containing 50 mM octopamine and 500 µM SPT-enol or DMSO (control) and transferred onto plants after 48 hours. The data are given in average ± SD (n=10 for nematode numbers and n=30 for female sizes).

**Supplemental Figure 3.**
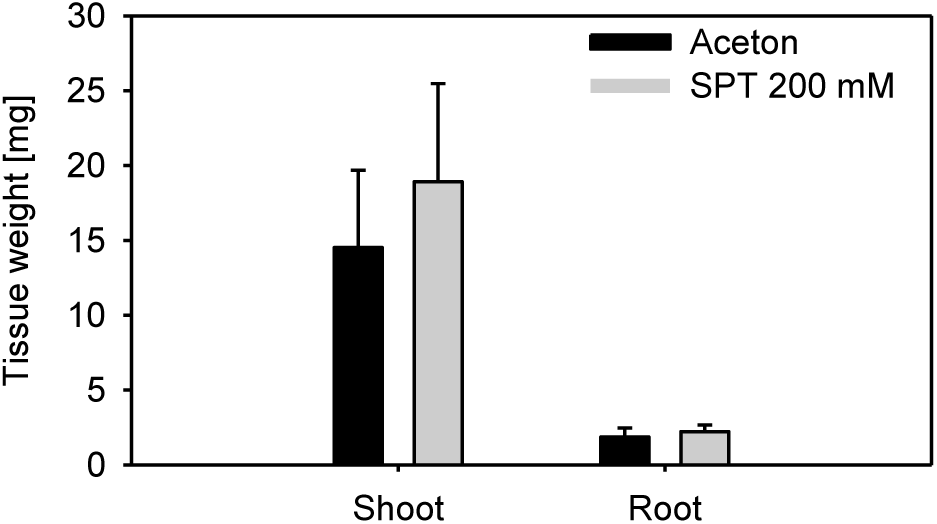
Effect of SPT on *A. thaliana* growth after foliar application. 3 µl of 200 mM SPT was applied onto the largest green leaf of 10-day-old *A. thaliana*. 10 days later, root and shoot weights were recorded (mg). No significant differences were found for root and shoot weight as compared to the control. The data are given in average ± SD (n=5).

**Supplemental Figure 4:**
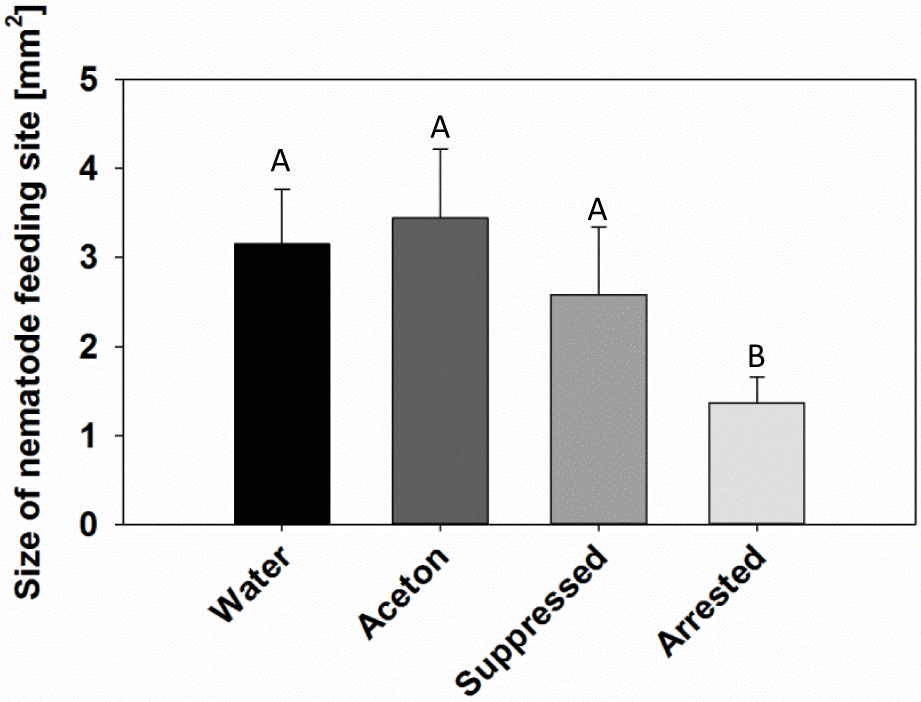
Effect of SPT on development of nematode feeding site. The sizes of nematode feeding sites were determined 14 DPI. The sizes of “suppressed” females are not significantly different as compared to the controls. Feeding sites of “arrested” nematodes are significantly smaller. The data are given in average ± SD. Anova P < 0.001, Tukey Test, N=30.

**Supplemental Figure 5:**
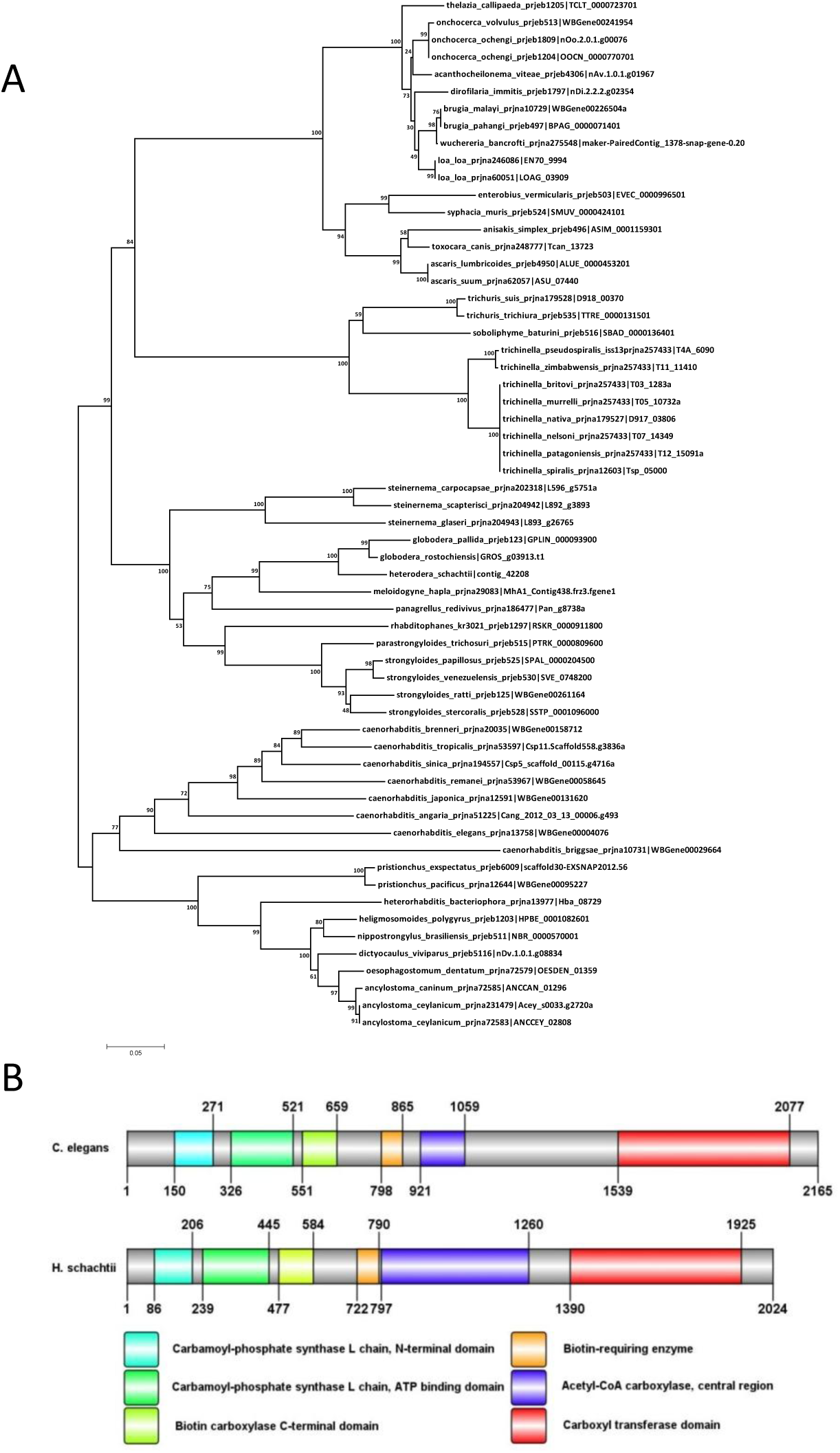
Sequence analysis of ACC from *H. schachtii*. (A) Phylogenetic tree of ACC protein sequences of nematodes obtained from public databases. The mRNA sequence of the *C. elegans* POD-2 was used as query to find the corresponding *H. schachtii* ortholog. We found an mRNA contiq that contains an ORF coding for a putative protein with 2024 amino acids. This protein contains all functional domains required for ACC activity and is similar to other ACC sequences found in public databases. Protein alignment shows that HsACC is most similar to putative ACC’s of the cyst nematodes *Globoderapallida and Globoderarostochiensis*. (B) Domain structure comparison of *C. elegans* and *H. schachtii* ACC. Domain structures were illustrated using DOG2.0 (Ren et al. 2009).

**Supplemental Figure 6:**
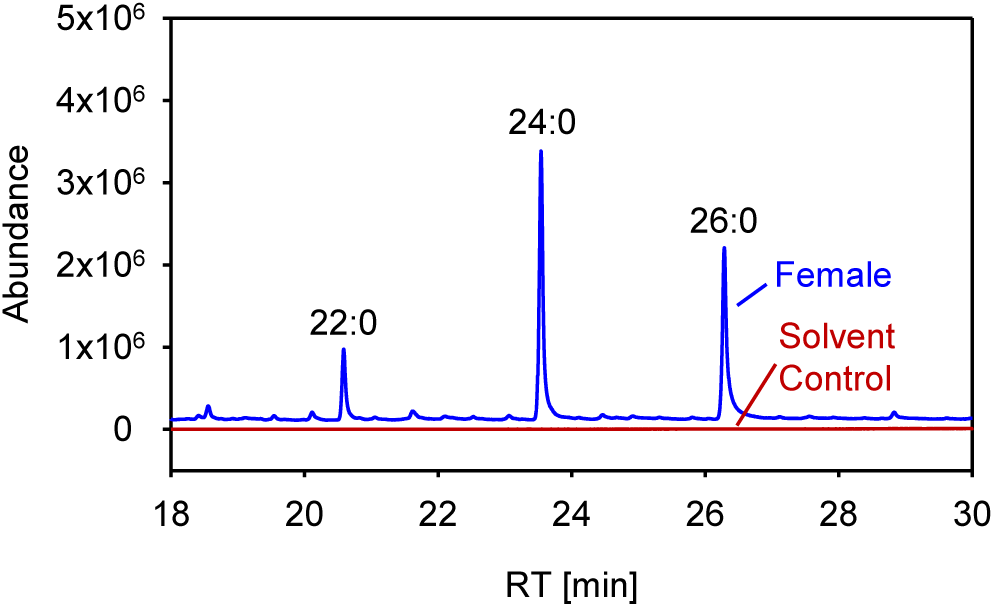
Chromatogram of VLCFA in surface coats of *H. schachtii* females. 14-day-old *H. schachtii* females were removed from *Arabidopsis* roots and incubated in chloroform for 1 min to remove the surface lipids. These surface lipids were transmethylated using 1 N HCl in MeOH and the resulting fatty acid methyl esters were analysed by GC-MS.

**Supplemental table 1.** Molecular species composition of abundant TAGs in *C. elegans*. The molecular species composition of TAGs was determined by scanning Q-TOF MS/MS spectra for neutral loss of fatty acyl-NH3. The fatty acid composition of abundant molecular species is shown.

| Triacylglycerol | [M+NH<sub>4</sub>]<sup>+</sup> | Molecular Species |
| --- | --- | --- |
| 50:3 | 846.6612 | 16:1-17:cyclo-17:cyclo |
| 51:3 | 860.6768 | 17:cyclo-17:cyclo-17:cyclo |
| 52:3 | 874.6925 | 17:cyclo-17:cyclo-18:1 |
| 53:3 | 888.7081 | 17:cyclo-18:1-18:1, 17:cyclo-17:cyclo-19:cyclo |
| 54:3 | 902.7238 | 18:1-18:1-18:1, 17:cyclo-18:1-19:cyclo |
| 55:3 | 916.7394 | 18:1-18:1-19:cyclo, 17:cyclo-19:cyclo-19:cyclo |

**Supplemental table 2.**
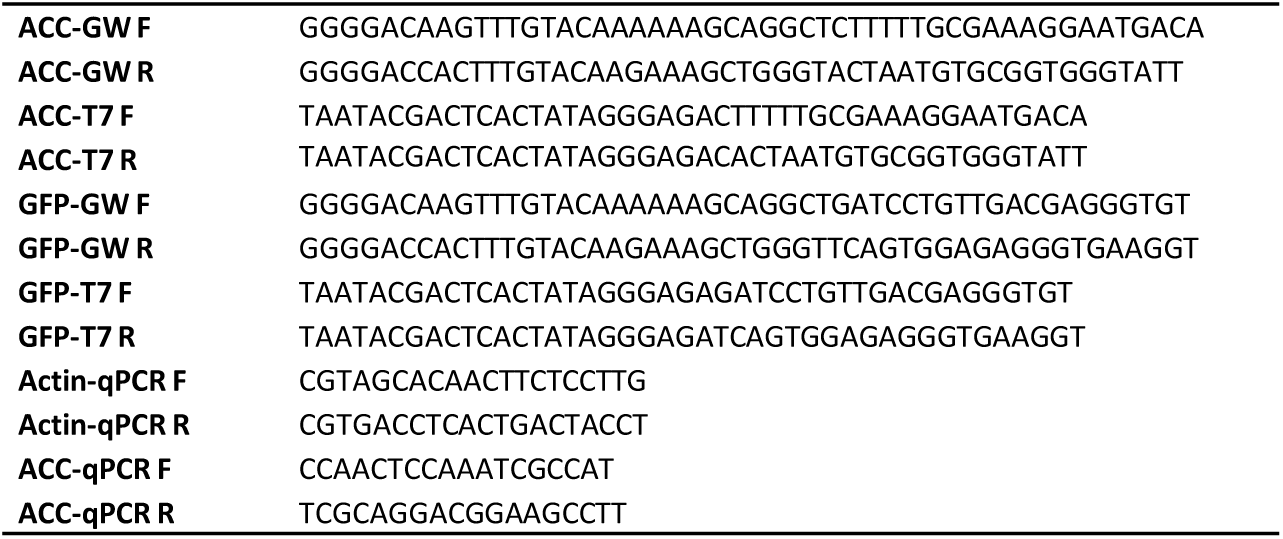
Oligonucleotides used in this study. Oligonucleotide names and sequences (5’-3’) are shown.

